# The hemolymph of *Biomphalaria* snail vectors of schistosomiasis supports a diverse microbiome

**DOI:** 10.1101/2020.04.22.056168

**Authors:** Frédéric D. Chevalier, Robbie Diaz, Marina McDew-White, Timothy JC. Anderson, Winka Le Clec’h

## Abstract

The microbiome – the microorganism community that is found on or within an organism’s body – is increasingly recognized to shape many aspects of its host biology and is a key determinant of health and disease. Microbiomes modulate the capacity of insect disease vectors (mosquitos, tsetse flies, sandflies) to transmit parasites and disease. We investigate the diversity and abundance of microorganisms within the hemolymph (*i.e.* blood) of *Biomphalaria* snails, the intermediate host for *Schistosoma mansoni*, using Illumina MiSeq sequencing of the bacterial 16S V4 rDNA. We sampled hemolymph from 5 snails from 6 different laboratory populations of *B. glabrata* and one population of *B. alexandrina*. We observed 279.84 ± 0.79 amplicon sequence variants (ASVs) per snail. There were significant differences in microbiome composition at the level of individual snails, snail populations and species. Snail microbiomes were dominated by Proteobacteria and Bacteroidetes while water microbiomes from snail tank were dominated by Actinobacteria. We investigated the absolute bacterial load using qPCR: hemolymph samples contained 2,784 ± 339 bacteria per μL. We speculate that the microbiome may represent a critical, but unexplored intermediary in the snail-schistosome interaction as hemolymph is in very close contact to the parasite at each step of its development.

## INTRODUCTION

The microbiome – the community of microorganisms that inhabit both the human body and that of multiple other phyla (Lee and Hase, 2014) – is increasingly recognized to play a critical role in many aspects of biology and as a key determinant of health and disease. For example, the human gut microbiome is thought to play a role in the development of diabetes, rheumatoid arthritis, muscular dystrophy, multiple sclerosis, and obesity (Kinross *et al.*, 2011). In invertebrates, symbiotic bacteria are involved in synthesizing essential amino acids or vitamins (Hosokawa *et al.*, 2010; Oliver *et al.*, 2010), in digesting cellulose (Brune, 2014), in protection against parasites (Oliver *et al.*, 2010), in feminization and parthenogenesis (Werren *et al.*, 2008), and even in speciation (Flintoft, 2013).

In host-parasite interactions, the host microbiome frequently plays an important role in the ability of parasite to colonize its host. This tripartite interaction (i.e. host-microbiome-parasite/pathogen) has been investigated widely in invertebrate hosts. Recent research on the widespread endosymbiotic bacteria *Wolbachia* demonstrates that *Wolbachia*-infected mosquitoes are unable to transmit dengue fever (Slatko *et al.*, 2014). This bacterium is now a major focus of efforts to control or eliminate dengue, and has stimulated new interest in utilizing bacterial symbionts to modify disease vectors (Slatko *et al.*, 2014). Similarly, the aphid microbiome protects these insects against parasitoid wasps (Oliver *et al.*, 2010), the microbiome of sandflies provides protection against *Leishmania* infection (Sant’Anna *et al.*, 2014), while the microbiome of mosquitoes and tsetse flies impedes infection by malaria parasites and trypanosomes respectively (Stathopoulos *et al.*, 2014).

Among human helminth parasites, schistosomes are the most important in terms of global morbidity and mortality, infecting over 206 million people in 78 countries in 2016 (WHO), and causing more than 200,000 deaths per year. Schistosome parasites have a complex lifecycle, involving freshwater snails as intermediate hosts and mammals as definitive hosts. In humans, adult schistosomes live in the blood vessels surrounding the intestine, and pathology results from immune reactions to eggs which become trapped in the liver, resulting in granulomas, calcification of blood vessels, portal hypertension and liver failure (van der Werf *et al.*, 2003; Colley *et al.*, 2014). There is no licensed vaccine, and control is reliant on a single monotherapy (praziquantel) to which resistance has been recorded, so there is an urgent need for alternative approaches to schistosome control. Understanding the snail-schistosome interaction is therefore critical to develop these approaches. When a single parasite larva (miracidium) penetrates a snail, this develops into a primary sporocyst, which then divides asexually to generate multiple clonally-produced secondary sporocysts in the snail tissue. These produce hundreds to tens of thousands of motile cercariae larvae that are released into the water where they penetrate and infect the definitive host (human or rodent). Snail populations vary in susceptibility to schistosomes (Files and Cram, 1949) and while genetics certainly plays a role (Tennessen *et al.*, 2015), the mechanisms determining whether infections develop are poorly understood and the influence of the microbiome on schistosome parasite development is currently unknown.

The *Biomphalaria* snail microbiome was firstly characterized using classical microbial culture methods and biochemical characterization almost 40 years ago (Ducklow *et al.*, 1979). Bacteria were found in the whole snail but also on its surface, gut and body cavity. With this relatively limited technique, the authors observed changes in whole snail microbiome when those were moved between different environments (Ducklow *et al.*, 1979). More recently, two papers (Allan *et al.*, 2018; Huot *et al.*, 2020) have examined *Biomphalaria* spp. microbiomes using deep sequencing of rDNA. Allan *et al.* (2018) showed that snails carrying alleles conferring resistance to *Schistosoma mansoni* infection, show distinctive microbiomes from those carrying sensitive alleles, suggesting possible links between the microbiome composition and *S. mansoni* susceptibility. Huot *et al.* (Huot *et al.*, 2020) examined core microbiomes from several laboratory populations, showing a high specificity between these microbiomes and their snail host populations, as well as complete congruence between core microbiome and host phylogeny. Importantly, both these studies examined microbiomes of whole snails, so it is unclear whether the bacteria characterized are from the gut, shell surface, mantle cavity or internal organs of the snail, each of which may have a distinctive microbiome. Rather than characterize composite microbiomes from whole snails, we focus on the microbiomes from a single tissue – the hemolymph (*i.e.*, invertebrate blood). We chose this tissue because developing schistosome parasites are in continuous contact with the snail hemolymph during the 30-day development within the snail. Therefore the hemolymph microbiome is likely to be most relevant for understanding interactions between snails, bacteria and developing schistosomes. Furthermore, evidence from other mollusk species provide a precedent for investigating the *Biomphalaria* microbiome: hemolymphs from several fresh and seawater mollusks and crustaceans are known to harbor specific microbiomes (Antunes *et al.*, 2010; Desriac *et al.*, 2014; Vezzulli *et al.*, 2018; Zhang *et al.*, 2018; Offret *et al.*, 2019; Musella *et al.*, 2020; Table 1) that can be modified by environmental stress (Lokmer and Mathias Wegner, 2015), pollutants (Leite *et al.*, 2017; Auguste *et al.*, 2019a) or pathogens (Vezzulli *et al.*, 2013; Lokmer and Mathias Wegner, 2015).

**Table 1.** Aquatic organisms with known hemolymph microbiome.

| Organism<br>(vernacular name) / phylum | Biotope | Hemolymph microbiome<br>characterization | References |
| --- | --- | --- | --- |
| *Crassostrea gigas* (Pacific oyster) / Mollusk | Sea water | 16S rRNA gene amplicon (V1-V2) pyrosequencing | Lokmer and Mathias Wegner, 2015 |
|  |  | 16S rRNA gene amplicon HiSeq sequencing | Vezzulli *et al.*, 2018 |
|  |  | 16S rRNA gene amplicon (V1-V3) sequencing | Pathirana *et al.*, 2019 |
|  |  | Bacterial culture / CFU* counting | Desriac *et al.*, 2014 |
| *Crassostrea angulata* (Portuguese oyster) / Mollusk | Sea water | 16S rRNA gene amplicon (V3-V4) HiSeq sequencing/bacterial culture | Zhang *et al.*, 2018<sup>1</sup> |
| *Mytilus galloprovincialis* (Mediterranean mussel) / Mollusk | Sea water | 16S rRNA gene amplicon (V4) Ion Torrent sequencing | Auguste *et al.*, 2019b |
|  |  | 16S rRNA gene amplicon HiSeq sequencing | Musella *et al.*, 2020 |
| *Mytilus edulis* (Blue mussel) / Mollusk | Sea water | Bacterial culture / CFU* counting | Desriac *et al.*, 2014 |
|  |  | 16S rRNA gene amplicon sequencing | Offret *et al.*, 2019 |
| *Anodonta cygnea* (Swan mussel) / Mollusk | Fresh water | Bacterial culture / CFU* counting | Antunes *et al.*, 2010 |
| *Pecten maximus* (Great scallop) / Mollusk | Sea water | Bacterial culture / CFU* counting | Desriac *et al.*, 2014 |
| *Tapes rhomboides* (Clam) / Mollusk | Sea water | Bacterial culture/ CFU* counting | Desriac *et al.*, 2014 |
| *Ruditapes philippinarum* (Manila clam) / Mollusk | Sea water | Bacterial culture/16S rRNA gene amplicon sequencing | Leite *et al.*, 2017 |
| *Haliotis tuberculata* (Abalone) /Mollusk | Sea water | 16S rRNA gene amplicon sequencing | Offret *et al.*, 2019 |
| *Scylla paramamosain* (Mud crab) / Arthropoda | Sea water | 16S rRNA gene amplicon (V3-V4) HiSeq sequencing/bacterial culture | Zhang *et al.*, 2018<sup>1</sup> |
| *Litopenaeus vannamei* (Whiteleg shrimp) / Arthropoda | Sea water | 16S rRNA gene amplicon (V3-V4) HiSeq sequencing/bacterial culture | Zhang *et al.*, 2018<sup>1</sup> |
| *Marsupenaeus japonicas* (Kuruma shrimp) / Arthropoda | Sea water | Bacterial culture/ CFU* counting | Wang *et al.*, 2014 |
| *Panulirus ornatus* (Spiny lobster) / Arthropoda | Sea water | 16S rRNA gene amplicon (V1-V3) pyrosequencing | Ooi *et al.*, 2019 |

<sup>1</sup>For additional references on hemolymph microbiome related studies, see **Table 1** from Zhang *et al.*, 2018.
\*CFU: Colony Forming Unit

In this study, we characterized the hemolymph microbiome of two *Biomphalaria spp*. freshwater snail species (i.e. *B. glabrata* and *B. alexandrina*), both vectors of *S. mansoni* using a 16S rDNA approach. We demonstrate that snail hemolymph harbors a diverse microbiome, and that different *Biomphalaria* species and populations harbor distinctive hemolymph microbiomes that are independent of the water environment. As the microbiome is critical in other host-parasite systems we speculate that it may represent a critical, but unexplored, intermediary in the snail-schistosome interaction.

## RESULTS

### 1) Library statistics

We sequenced microbiomes from snail hemolymph and water samples from seven laboratory populations (Table 2). For each population, we selected 5 snails, efficiently removed shell contaminants by cleaning the shell surface with ethanol (Supp. Fig. 1) and collected hemolymph by heart puncture to avoid contamination from the tegument. In parallel, we collected a sample of the tank water the snails were collected from. To test the reproducibility of our protocol, we split each sample and prepared libraries independently for each half (Fig. 1). After MiSeq sequencing, we obtained an average of 155,491.5 ± 29,277.9 reads per library (mean ± s.d.) (Supp. Table 1). We retained on average 87 ± 0.04% of the initial reads after sequencing error filtering, 85 ± 0.03% of the initial reads after denoising the data using the dada2 module, 73 ± 0.05% of the initial reads after merging reads and 71 ± 0.06% of the initial reads after removing chimeras. Sequencing depth was relatively even between sample types and replicates (Supp. Table 1). Rarefaction curve analysis of each library showed a plateau, indicating that our sequencing effectively captured the microbiome diversity (Supp. Fig. 2). Taxonomy assignment of the 16S rDNA sequences from all libraries revealed a total of 2,688 amplicon sequence variants (ASVs). Among them, 897 had unassigned taxonomy as typically seen in microbiome study of non-model organisms, with 39 (4.3%) being eukaryote contaminants (snails or others), 836 with match to uncultured bacteria mainly, and 22 (2.5%) with no similarity to known sequences suggesting potential new bacterial or archaeal species. The eukaryote contaminants and 5 ASVs with mitochondria or chloroplast assignment were removed in subsequent analysis.

**Table 2.** Snail laboratory populations used in this study and their characteristics.

| ***Biomphalaria***<br>**species** | ***Biomphalaria***<br>**laboratory population** | **Origin (Country,**<br>**town, snail cross)** | **Albinism** | **Inbreeding** | **References** |
| --- | --- | --- | --- | --- | --- |
| *B. alexandrina* | Ba | Egypt | No | No | Theodor Bilharz Research Institute, Egypt |
| *B. glabrata* | BgBS90 | Brazil | No | No | Paraense and Correa, 1963 |
|  | BgBRE | Brazil (Recife) | Yes | No | Theron *et al.*, 2014 |
|  | BgNMRI | Puerto Rico | Yes | No | Lewis *et al.*, 1986 |
|  | Bg26 | BgM line derived<br>(cross between a<br>Brazilian and a Puerto<br>Rican snail) | Yes | Yes | Bonner *et al.*, 2012 |
|  | Bg36 |  | Yes | Yes |  |
|  | Bg121 |  | Yes | Yes |  |

**Fig. 1.**
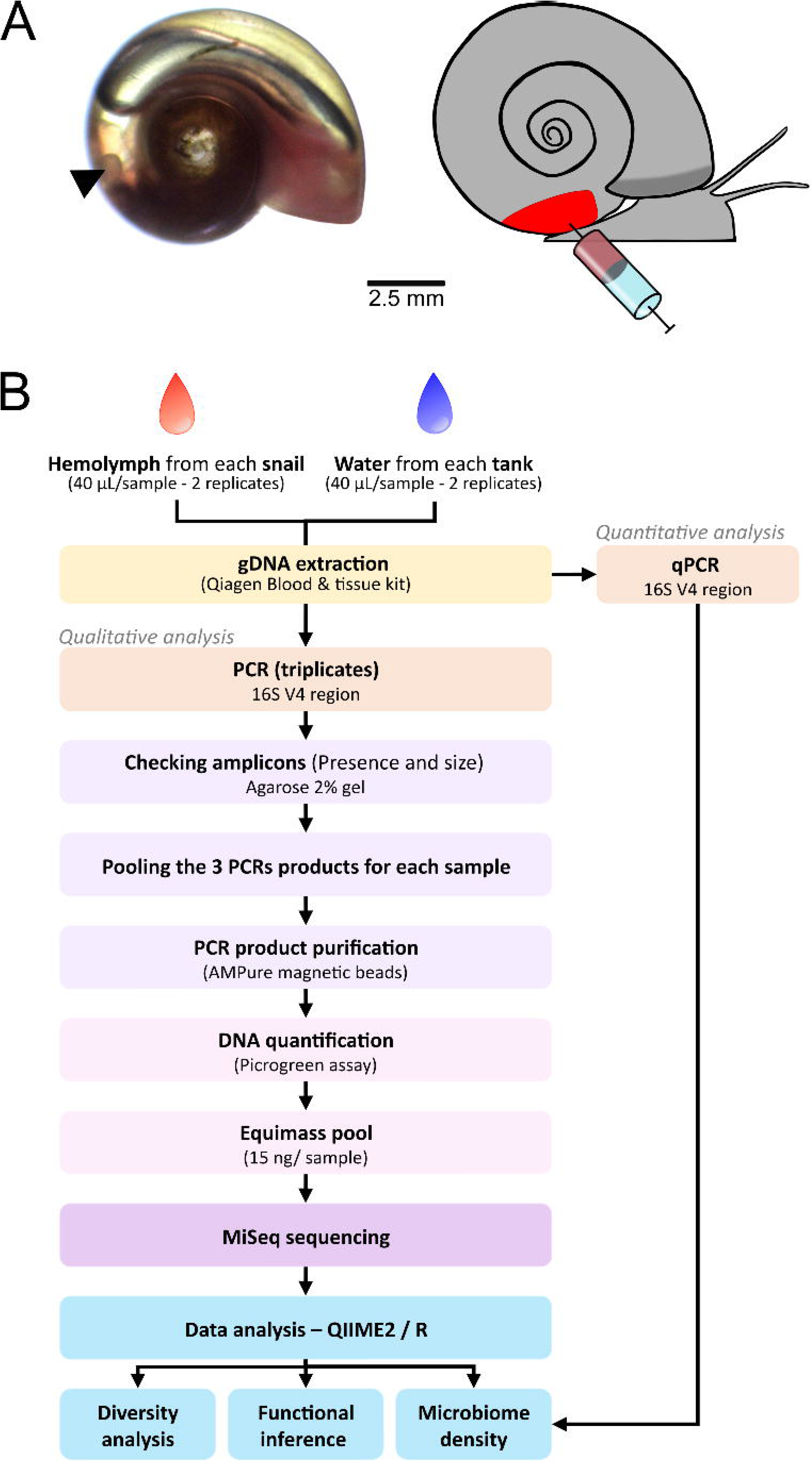
Site of hemolymph sampling and workflow of the library preparation and the analysis. **A.** Snail hemolymph was sampled directly by heart puncture using a syringe. Left panel shows the position of the heart (arrow) on an albino snail. Right panel shows a schematic representation of the puncture. **B.** Work flow of the sample preparation and data processing.

### 2) Microbiome diversity

#### a) α diversity (microbial diversity within samples)

We computed several α diversity indices (Table 3) to investigate the bacterial richness (the number of taxa) and diversity (a composite measure of richness and abundance of bacterial taxa) within hemolymph and water samples (Fig. 2). The observed species richness and the Chao1 index revealed that snail hemolymph had higher bacterial richness than water, with the exception of the BgNMRI population. Overall, we found twice the number of ASVs in snail hemolymph (279.84 ± 0.79; mean ± s.e.) than in the tank water (145.43 ± 4.02). Most of the snail populations showed similar mean richness, with Bg26 showing the highest and BgNMRI the lowest. The close similarity between the observed richness and the Chao1 index plots suggests that we effectively captured rare taxa, consistent with the rarefaction analysis.

**Table 3.** Description of α diversity indices used.

| Index | Description |
| --- | --- |
| Observed richness | Raw number of ASVs |
| Chao1 richness | Estimation of the true richness by adjusting the raw number of ASVs for rare taxa |
| Faith's Phylogenetic diversity index | Richness index based on phylogenetic distance between ASVs |
| Shannon index | Measure of heterogeneity in taxa representation (evenness × richness) |
| Simpson evenness index | Measure of true evenness of taxa within a sample independently of taxa richness. A value close to 0 means that the taxa are unevenly distributed in a sample. |
| Simpson dominance index | Measure of taxa diversity within a sample. This index is weighted towards the most abundant taxa and influenced by taxa richness. A value close to 0 means that the sample is dominated by few taxa. |

**Fig. 2.**
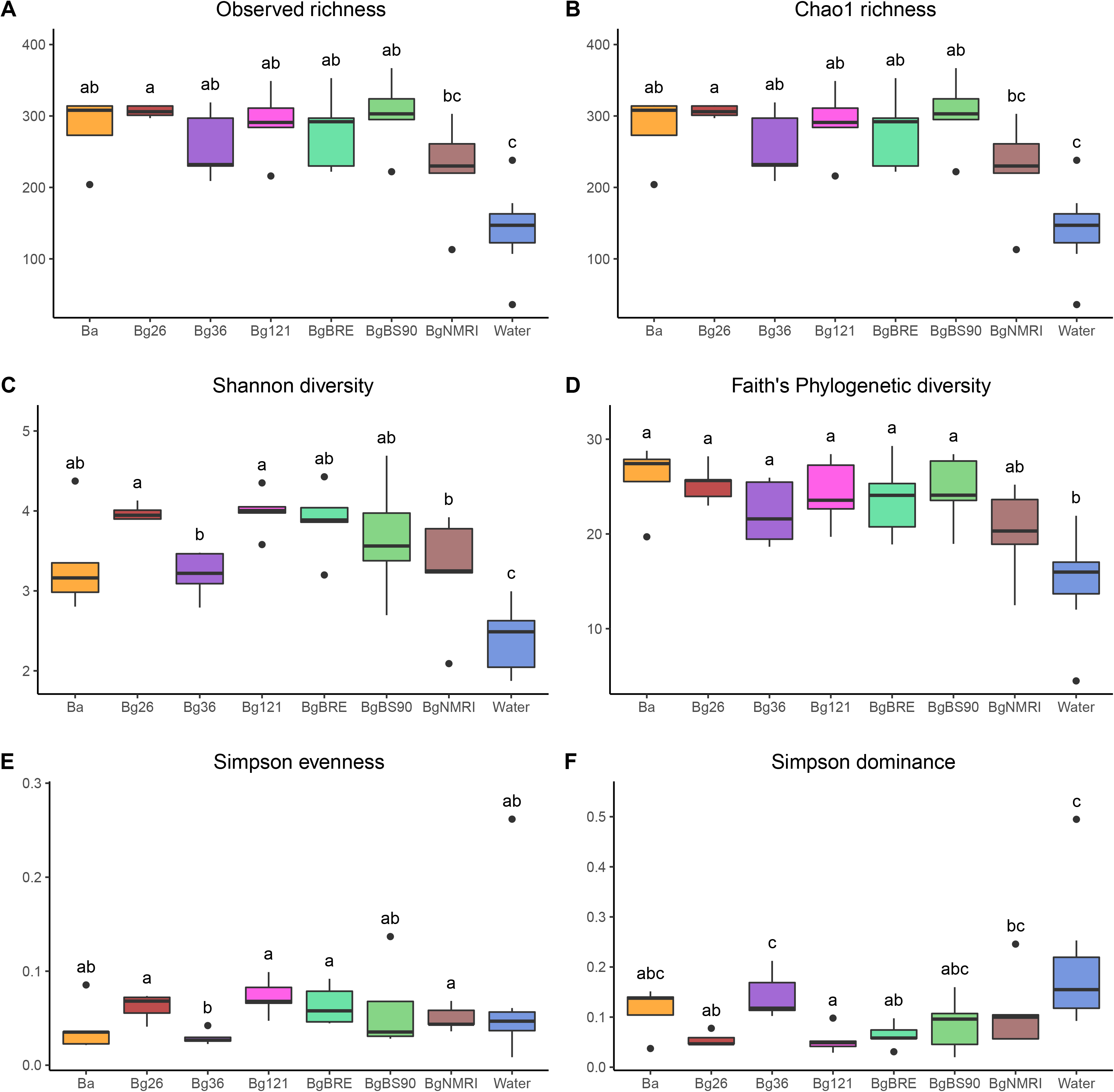
α diversity indices. Six different alpha diversity indices were computed (see Table 2 for index description) revealing that snail hemolymph harbors bacteria communities and those communities are richer and more diverse that those of the water. Variation in diversity exists between snail populations. All these bacterial communities are dominated by characteristic bacterial species. Each box plot represents the median and the interquartile ranges of the distribution. Groups (i.e. snail populations or tank water) not connected by the same letter are significantly different (post-hoc test).

The bacterial richness of the hemolymph is correlated with a higher bacterial diversity as expected. The Shannon diversity index showed lower mean diversity in the water when compared to the hemolymph. Among the snail populations, we observed the highest mean diversity in Bg26 and Bg121 and the lowest in Bg36 and BgNMRI. Individuals from the BgBS90 population showed the greatest variability between snails suggesting high heterogeneity in this population. The Faith’s phylogenetic diversity showed a similar trend: we observed a lower diversity in the water while hemolymph displayed similar diversity across snail populations. This suggests that the differences between snail populations observed with the Shannon index were likely due to differences in evenness rather than in richness.

The Simpson indices for evenness and dominance were low for both hemolymph and water samples, in accordance with the over representation of particular taxa. Hemolymph from Bg36 population, closely followed by Ba population, showed the lowest evenness and the highest dominance of the snail populations. BgNMRI snails, which had similar richness and diversity profile as Bg36 snails, showed slightly more evenness in their bacterial community. Water samples, which consistently showed low richness and diversity, showed a comparable evenness to snail samples but a higher dominance than some of the snail populations consistent with the Shannon diversity index.

Data from the replicated libraries showed similar results. We obtained a strong correlation (0.85 < R^2^ < 0.98, p < 0.01) between the replicated libraries (Supp. Fig. 3) when comparing the different indices, highlighting the robustness and reproducibility of our protocol.

#### b) β diversity (microbial diversity between samples)

We computed several β diversity indices to investigate how the bacterial communities discriminate between each snail population (Fig. 3, Supp. Fig. 4). We first compared microbiomes using qualitative metrics that are based on the presence or absence of bacterial taxa (Jaccard index) adjusted with phylogenetic distance (unweighted unifrac index), independently of taxa abundance. We statistically compare these metrics using Permanova analysis. These analyses demonstrate that water samples have significantly different bacterial communities from the snail populations. Similarly, all pairwise comparisons between snail communities showed statistically different composition. Comparisons of centroid distances revealed a significantly greater distance between Ba and Bg than between each Bg populations in at least one of our replicate for both indices (Jaccard index: Ba-Bg distance = 0.62, Bg populations distance = 0.57, p = 0.021; unweighted unifrac index: Ba-Bg distance = 0.49, Bg populations distance = 0.45, p = 0.010). In addition, the principal component analysis from the Jaccard index and, to a lesser extent, from the unweighted unifrac revealed clear groups: (i) Ba is distant from all Bg snail populations as supported by centroid distances strongly suggesting that hemolymph microbiome can discriminate snail species; (ii) Two clusters of Bg snails can be identified: the first cluster comprises BgBS90, BgBRE, Bg121 and Bg36 (all from Brazil) and the second cluster comprises Bg26 and BgNMRI (from Carribbean) (Fig. 3A-B).

**Fig. 3.**
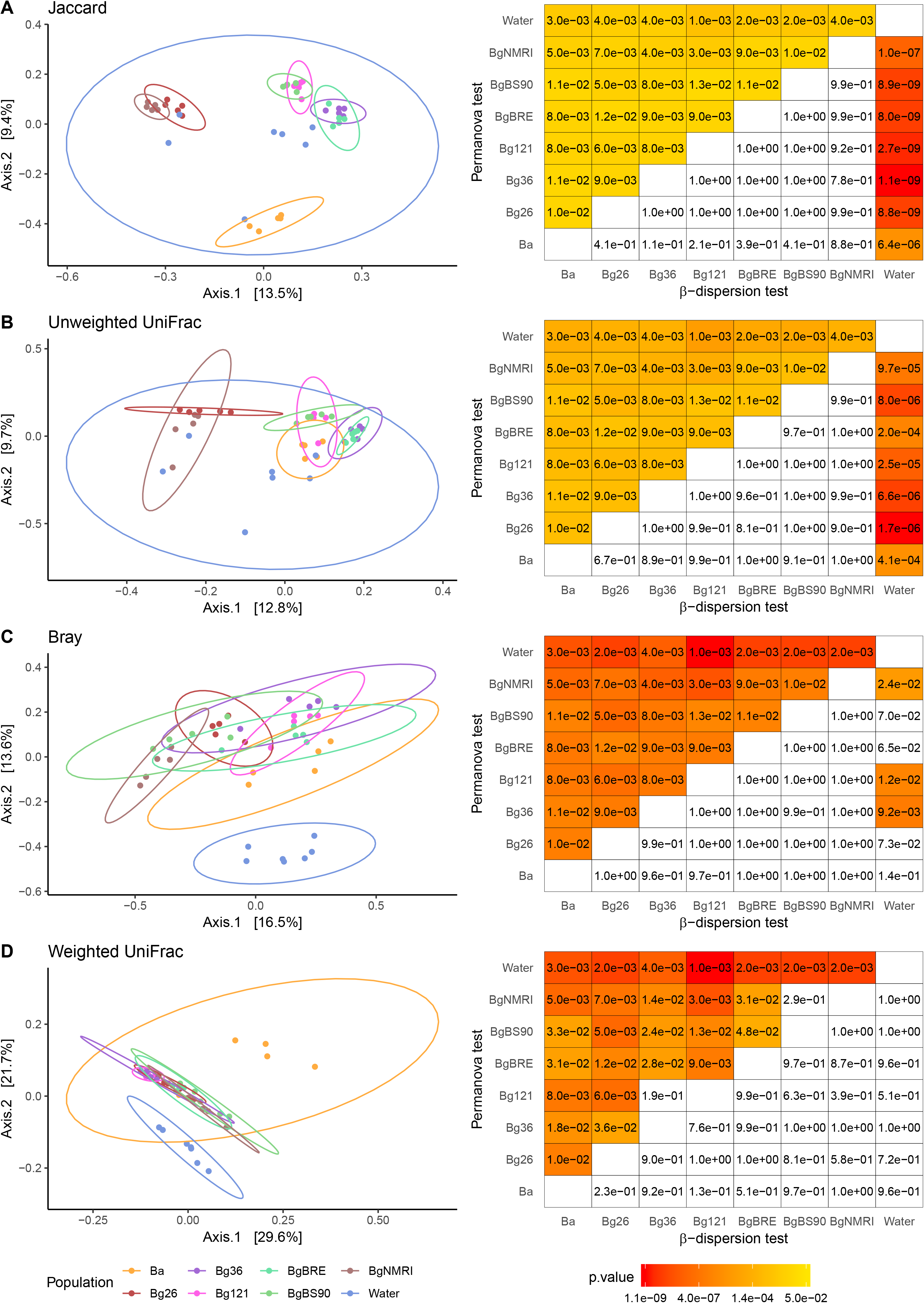
β diversity indices. For each index (A. Jaccard, B. Unweighted unifrac, C. Bray, D. Weighted unifrac), the first two axes of the principal coordinates analysis (PCoA) are represented. The two first indices represent a qualitative measure (absence or presence of bacterial taxa) while the two last are quantitative (adjusted with taxa abundance). The matrices on the right side of each index showed the significance level of the Permanova test (differences between populations) and the β dispersion test (homogeneity of variance between populations). All four indices showed that water microbiome was different from snail hemolymph microbiomes, that the two species of snail (*B. glabrata* and *B. alexandrina*) can be differentiated on a quantitative and qualitative approaches, and that *B. glabrata* populations were differentiated based on rare taxa (qualitative aspect) and this differentiation could be linked to their geographic origin. The ellipse represents the multivariate normal distribution.

We also compared populations of snails or water microbiome for the Jaccard and unweighted unifrac indices using a β dispersion test (Fig. 3A-B). This test compares the homogeneity of β-diversity measures across different sampling units (individual snails, or water from different tanks) between populations. All the snail populations showed similar variance when compared to each other. In contrast, the water population showed over dispersion when compared to all the snail populations. This may be explained by the fact that samples of water come from different aquariums, each of them carrying characteristic bacteria, while individual snails within a population come from the same tank.

We performed similar comparisons using quantitative metrics that take into account the abundance of each bacterial taxa (Bray index) adjusted with phylogenetic information (weighted unifrac index). Comparisons of Bray indices using a permanova analysis revealed that all snail populations and water showed different composition from each other (Fig. 3C). We obtained similar results when adding phylogenetic information (Fig. 3D) except for the pairs Bg121-Bg36 and BgBS90-BgNMRI, suggesting that these microbiomes shared similar abundant taxa between these pairs (Fig. 3D). Centroid distances were again a greater between Ba and Bg than between each Bg populations in at least one of our replicate for both indices (Bray index: Ba-Bg distance = 0.69, Bg populations distance = 0.63, p = 0.014; weighted unifrac index: Ba-Bg distance = 0.26, Bg populations distance = 0.18, p = 3.7 × 10^−5^). These results suggest that abundant bacterial taxa drive the distinction between species and within populations and revealed similar microbiomes in some genetically related snail populations (Bg121-Bg36).

#### c) Taxonomic diversity

We found dramatic differences between populations in terms of taxonomic composition of bacteria (Fig. 4). While considering the populations, hemolymph microbiomes were dominated by bacteria from the phyla Proteobacteria (in all Bg populations) and Bacteroidetes (in Ba population) while water microbiomes were dominated by bacteria from the Actinobacteria phylum (Fig. 4C). These differences in dominating phyla reflect the previous observations from the β diversity analyses: a clear distinction (i) between the water and snails and (ii) between the two snail species (i.e., Bg and Ba). We also observed differences in proportion of bacteria from minor phyla among the different snail populations. For instance, Bg26 showed a greater proportion of the phylum Acidobacteria in most of the comparisons while BgNMRI and Ba showed the lowest proportion of the phylum Verrucomicrobia in most of the comparisons (Supp. Table 2). These differences may reflect differences in the physiology of these different populations. Microbiomes from snail samples (and at a lesser extent from water samples) also contained a relatively high proportion of bacteria with unassigned taxa: 5% to 43% of ASVs from snails while 0.2% to 17% of ASVs from the water were uncharacterized.

**Fig. 4.**
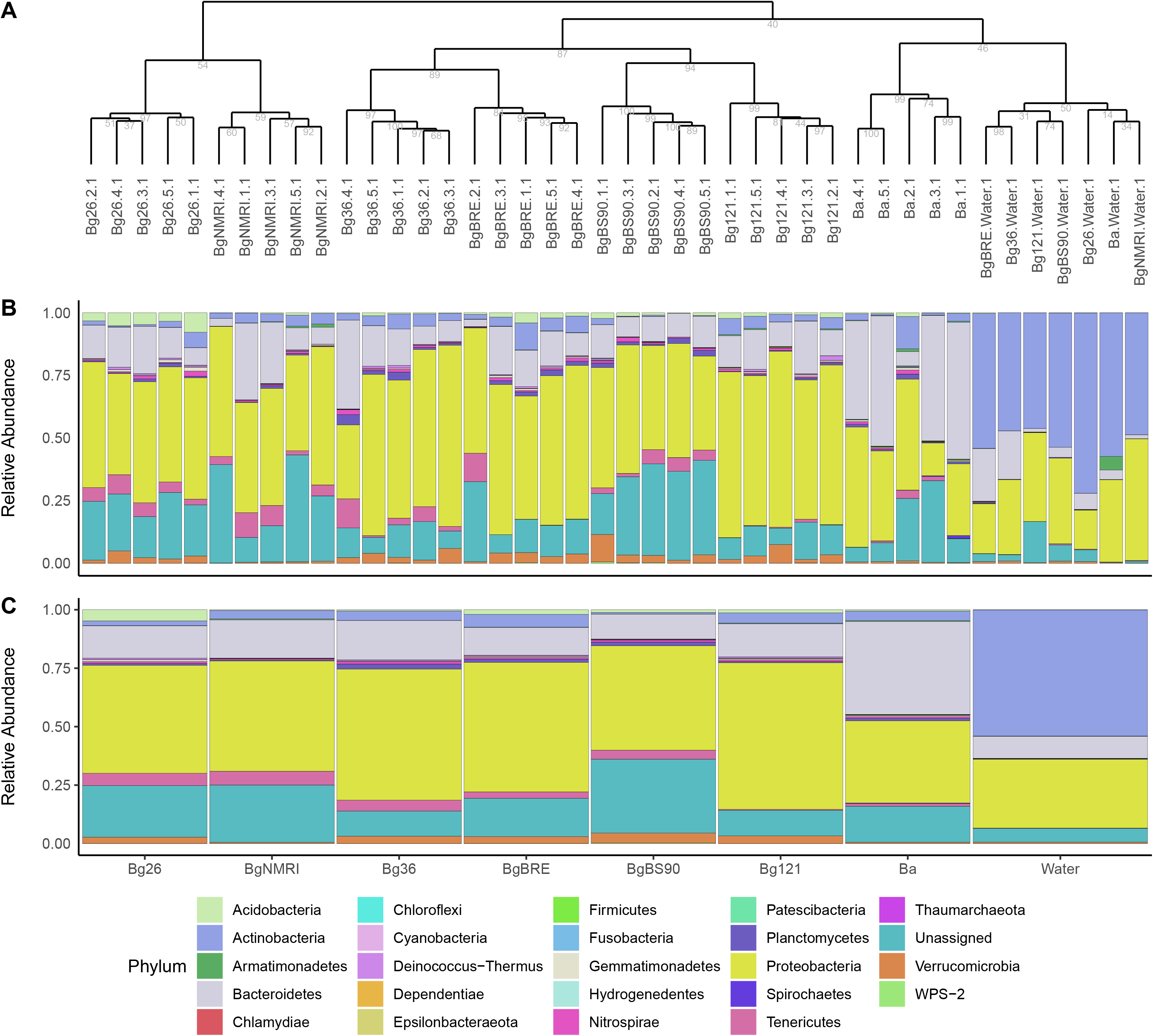
Taxonomic diversity in each sample and populations. **A.** Dendrogram representing the distance between samples based on their microbiome composition. Number under each node represents bootstrap values from 1,000 iterations. Samples from the same population cluster together and links between populations reflects results obtained from the beta-diversity analysis. **B and C**. Relative abundance of the different phyla present in each samples from the dendrogram (B) and in each corresponding populations (C). Water microbiome showed dramatic differences when compared to snail hemolymph microbiomes. Snail microbiomes showed also specificities between populations.

Individual snails from a population tended to show a similar taxonomic composition as they clustered together (Fig. 4A). The relatedness between populations reflects what is observed with the qualitative analysis of the β diversity (Fig. 3A-B).

We identified 93 snail and population specific ASVs not found in water and 1 water specific ASV not found in snails (Supp. Fig. 5). The count of these ASVs ranged from 2 to 36,770 with a prevalence ranging from 1.19 to 73.81%. The Ba population showed most of the specific ASVs. Bg121, BgBRE, BgBS90 and BgNMRI showed the lowest number of ASVs and Bg121, BgBRE, BgBS90 showed very low prevalence of these ASVs. These snail specific ASVs suggest that they may be specifically adapted to their hosts.

We qualitatively compared our microbiome compositions to the main bacterial genus or family found in other whole snail microbiome studies (Ducklow *et al.*, 1979, 1981; Allan *et al.*, 2018; Huot *et al.*, 2020). We found bacteria from the *Pseudomonas, Acinetobacter, Aeromonas* genera and the Enterobacteriaceae family identified in Caribbean field and laboratory populations (Ducklow *et al.*, 1979) in at least one individual of each population, with the exception of Acitenobacter, which as absent from all BgNMRI snails. However, we did not find bacteria from the *Vibrio, Pasteurella*, and *Moraxella* genera found in by Ducklow *et al.* (1979, 1981). We also found bacteria from the *Gemmatimonas* genus in at least one individual snail from each population. This genus was previously associated with BgBS90 snail genotypes showing resistance to schistosome infection (Allan *et al.*, 2018). We did not find the *Micavibrio, Parabacteroides* and *Halobacteriovorax* genus identified by Allan *et al.* (2018). We found bacteria from the *Pseudomonas, Pirellula, Candidatus, Mesorhizobium* genus and the Flavobacteriaceae family previously identified in *Biomphalaria* snails (Huot *et al.*, 2020) but we did not find bacteria from the *Mycoplasma, Planctomyces* and *Odyssella* genus found in the same study. These absence of these bacterial taxa from our data set may reflect that these genera are in low concentration or truly absent from the hemolymph, or that differences in the protocols (16S region amplified, primers, database, pipeline used) may have precluded the identification from our current dataset.

Finally, the reproducibility of our protocol is again confirmed in this taxonomic analysis (Supp. Fig. 6). All the replicated libraries clustered together, with the exception of one BgNMRI sample.

### 3) Microbiome density

We investigated the bacterial density by performing real time quantitative PCR on our samples, and corrected the results to estimate the number of bacteria (Fig. 5A-B). We quantified an average of 4,490 16S rDNA copies per μL of sample (range: 186 to 59,487). We found the highest average copy number in BgNMRI and the lowest in BgBRE and water (Fig. 5A) (Kruskal-Wallis test: Χ^2^=26.494, df=7, p=0.0004108 followed by post-hoc pairwise comparison test).

**Fig. 5.**
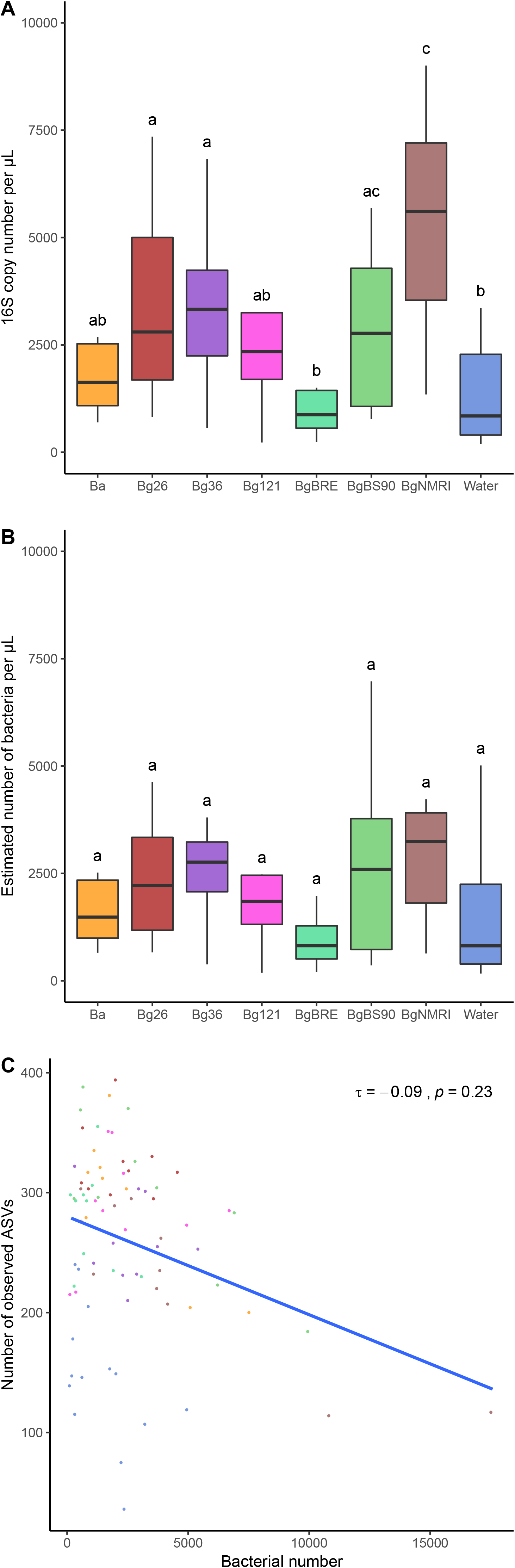
Comparison of microbial density and diversity. **A.** Raw 16S copy number per microliter of hemolymph and water sample quantified using qPCR. **B.** Estimation of bacteria per microliter after adjusting 16S copy number using PiCRUST2 input. **C.** Correlation between observed number of ASVs and estimated bacteria number. The number of bacteria present in a sample does not correlate with the number of ASVs observed. The dot colors are consistent with the population colors in the above panels. The different letters on panel A and B indicate significant differences in density means. Groups (i.e. snail populations or tank water) not connected by the same letter are significantly different (pairwise comparison post-hoc test, p < 0.05).

Bacteria usually carry more than one 16S gene copy in their genome (Větrovský and Baldrian, 2013). We corrected the raw density data based on 16S copy number using the estimation of 16S copy number per ASV computed by PiCRUST2. This normalization gave an average of 2,784 ± 339 (mean ± s.e.) (range: 186 to 17,578) estimated bacteria per μL in snail samples and 1,475 ± 382 in water samples (range: 169 to 5,014) (Fig. 5B). This did not change drastically the trend already observed with the raw copy number quantification. However, normalization narrowed the global and per population variation and removed any significant differences between populations (Kruskal-Wallis test: Χ^2^=16.22, df=7, p=0.023). When considering an average volume of 100 μL of hemolymph per adult snail (personal observation) and an average of 2,784 estimated bacteria per μL in snail individuals, the snail hemolymph should be inhabited by an average of 278,400 bacteria (range: 18,600 to 1,757,800).

We examined the correlation between the estimated bacteria density and the observed microbial richness diversity. We found no correlation (Kendall’s τ=−0.076, p-value=0.3). This confirms that we can safely estimate microbial diversity without taking in account bacterial density.

### 4) Functional inference of the hemolymph microbiome

The microbiome in hemolymph could have a functional role in the snail physiology, just as the gut microbiome does in humans (Valdes *et al.*, 2018). We investigated the presence of particular metabolic pathways in snails and water samples based on 16S taxonomy using the PiCRUST2 pipeline. We identified 36 and 4 pathways overrepresented in hemolymph and water samples respectively, which is supported by our two technical replicates (Supp. Table 3). The 36 overrepresented pathways in hemolymph were involved in synthesis of vitamins/electron transporters, in degradation of sugar, amines and aromatic compounds, in nucleotide metabolism and in fermentation.

The pathways overrepresented in water samples were involved in synthesis of molecules related to cell wall, DNA synthesis, and detoxification. At least two out of the four overrepresented pathways are found in bacteria from the Actinobacteria phylum, the dominant phylum found in water (Fig. 4).

In both snail and water samples, other differentially represented pathways were found in only one of the two technical replicates. This could be due to differences in taxa representation and abundance and highlight the sensitivity of metabolic inference to this factor.

## DISCUSSION

### 1. Snail hemolymph supports a diverse microbiome

We have unambiguously shown the presence of diverse microbiomes in the snail hemolymph of one population of *Biomphalaria alexandrina* and six populations of *B. glabrata*. The snail microbiome contains hemocytes that protect against incoming microbes, so the existence of a flourishing microbiome in the hemolymph might seem surprising to some. Three lines of argument suggest that the hemolymph microbiomes we have described are real, rather than an artifact of our collection procedure:

i. The shell cleaning step of our protocol efficiently removed any traces of bacteria on the shell, ensuring no contamination from the outside environment during the heart puncture (see Supp. Fig. 1 and section 1 of Experimental procedures).
ii. The bacterial diversity of the snail microbiomes is distinctive from that of the water in the tanks. The hemolymph and water microbiome showed dramatic differences in dominating phyla (Proteobacteria (for all the Bg populations), Bacteroidetes (for the Ba population), and Actinobacteria phylum in the water microbiomes). The hemolymph microbiomes were more diverse than the water microbiomes using multiple different diversity metrics. Finally, different species or populations of snails show characteristic bacterial composition in their hemolymph. The hemolymph, and to a lesser extent, the water samples harbor uncharacterized taxa: a third of the ASVs identified did not have a taxonomic assignment.
iii. Bacteria were abundant in the hemolymph. We observed an average of 2,784 bacteria per microliter of snail hemolymph, with a maximum of 17,578 in the BgNMRI population. Assuming 100μL hemolymph per snail this equates to 278,400 bacteria, or 1,757,800 for BgNMRI.

Our demonstration that the *Biomphalaria* hemolymph contains a rich microbiome is consistent with published reports detailing the microbiome from other mollusks, crustaceans (Braquart-Varnier *et al.*, 2008; Desriac *et al.*, 2014; Wang and Wang, 2015; Vezzulli *et al.*, 2018; Table 1) and also from insects (Blow and Douglas, 2019). The intimate relationship between these organisms and their water environment and their semi-open vascular system may prevent them from having a sterile hemolymph. Instead, they may use a tolerance strategy as observed in the human gut (Swiatczak and Cohen, 2015). In insects, the presence of both transient and obligate bacteria in the hemolymph is now recognized (Blow and Douglas, 2019) but only few bacterial taxa can survive in such an environment. In contrast, our results show that the hemolymph of *Biomphalaria* snails is inhabited by a highly diverse microbiome, suggesting that the snail hemolymph environment is more permissive.

### 2. Do genetics or the environment drive microbiome differences?

Differences in microbial diversity between hemolymph samples reflected their species or populations of origin. At the species level, *B. glabrata* and *B. alexandrina* always clustered apart based on both qualitative and quantitative β diversity analysis. This was driven by the differences in bacterial taxa and their abundance between the two species despite that all these snails were exposed to the same food and the same water source at the time of the tank setup. Therefore, the differences in the microbial composition are likely to be genetically controlled, reflecting the evolutionary distance between the two species. At the population level, we could discriminate *B. glabrata* populations based on rare taxa (qualitative β diversity analysis). Interestingly, the distance between population clusters was consistent with geographical origins: Brazilian populations (BgBRE, BgBS90) clustered apart from the Caribbean population (BgNMRI), while inbred subpopulations from a cross between snails of Caribbean and Brazilian origin (Bonner *et al.*, 2012) tended to cluster with the Caribbean population (Bg26) or the Brazilian population (Bg36, Bg121). If genetics is the main driver, this would suggest that microbiome selection at the population level is different from selection at the species level. However, a strong influence of the environment could also explain this result: we sampled snails of each population from single tanks. The rare taxa distinguishing microbiomes from different snail populations could also be a signature of the tank microbiome.

An association between snail genetics and the composition of whole snail microbiome has been observed previously (Allan *et al.*, 2018). In this particular case, specific genotypes at a locus involved in snail compatibility with schistosome infection were linked to changes in two particular taxa (*Gemmatimonas aurantiaca* and *Micavibrio aeruginosavorus*). This locus likely contains immune genes that could specifically shape the whole snail microbiome. In addition, a recent analysis of whole snail microbiomes that included several Biomphalaria species and *B. glabrata* populations is consistent with our observation regarding species differentiation based on microbiome but failed to support *B. glabrata* population differentiation (Huot *et al.*, 2020). Three factors from the Huot *et al.* (Huot *et al.*, 2020) study could explain this difference: (i) the shell was removed prior to DNA extraction: this may have resulted in significant loss of hemolymph, reducing the contribution of hemolymph to the results, or (ii) their analysis focused on the core microbiome: excluding rare taxa may have limited their ability to detect population differentiation, or (iii) the composite microbiome from whole snails (including microbiomes from multiple different organs) was analyzed.

Previous studies on whole snail microbiomes using older, culture-based methodology showed a strong influence of the environment: translocating *B. glabrata* from one river to another had a dramatic impact on abundance of cultivable bacteria (Ducklow *et al.*, 1981). This change in microbiome could be due to uptake of new bacteria from this new environment or could result from the stress of the translocation leading to the modification of the immune system and consequently of the microbiome composition. To better understand the relative role of genes and environment in controlling microbiome composition, classical genetic crosses, and common garden rearing experiments will be needed.

### 3. Origin of the hemolymph microbiome

Microorganisms can be acquired directly from the environment or transmitted vertically from the parents. We observed shared bacterial taxa between snail hemolymph and water suggesting that exchanges between the environment and the snail is likely to occur. Exchange could take place at two main sites: the mantle cavity and the gut. The mantle cavity is likely involved in gas exchange between the hemolymph and the water (Jurberg *et al.*, 1997). Pinocytosis at the mantle epithelium and loose vascular connective under it (Sullivan and Cheng, 1974) could promote active or passive acquisition of microorganisms. The microsporidium *Capsaspora owczarzaki* was found in the mantle and pericardial explants strongly suggesting that microorganisms can pass through the mantle cavity (Stibbs *et al.*, 1979; Hertel *et al.*, 2002). The gut and its surrounding hepatopancreas where active phagocytosis happens (Van Weel, 1961) could be another site of microorganism acquisition. Amoebae (Richards, 1968) and microsporidia (Richards and Sheffield, 1970) can infect the gut and hepatopancreas suggesting that the complex gut microbiome (Van Horn *et al.*, 2012) could colonize the hemolymph through this route. This could be validated by comparing gut and hemolymph microbiome.

A second route of microbiome acquisition is a vertical transmission from parents to offspring. Microorganisms could be selectively added to the egg yolk or the egg mass fluid (i.e. fluid surrounding the eggs inside the egg mass) and could be integrated into the developing organism. Antimicrobial proteins have been identified in both egg yolk (Baron *et al.*, 2013) and egg mass fluid (Hathaway *et al.*, 2010) which, in addition to protecting the eggs from outside pathogens, could control the development of the transmitted microbiome. Vertical transmission of microorganisms via the egg cytoplasm (Werren *et al.*, 2008) or after spreading them on the egg surface (Abe *et al.*, 1995; Prado *et al.*, 2006) is common in invertebrates.

### 4. Microbiome functionality: transient passengers or co-evolved relationship

The primary role of the hemolymph is to distribute nutrients to organs and defend the snail against infections. The presence of a microbiome could result from a tolerance strategy to mitigate the high costs of maintaining a sterile hemolymph and could be comprised of commensal microorganisms only. However, snails could have evolved in order to take advantage of the presence of this microbiome. Hemolymph microbiomes could be involved in providing nutrients to their snail host. Pathway inference analysis from our microbiome data showed that hemolymph microbiome is enriched in pathways related to vitamin production. Vitamins are known to be essential in egg production: snails raised in axenic conditions were not able to lay eggs except with a supplement of vitamin E (Vieira, 1967). We did not find pathways involved in the production of this particular vitamin but for others (B6, B12, K12) which may also have a role in snail physiology. The vitamins B6 (pyridoxine), B12 (cobalamin), and K12 (menaquinol) are essential to animals and could be released into the hemolymph by the bacteria and absorbed by the snail tissues. We have also shown enrichment for other pathways like sugar and polyamine metabolism which could serve only bacteria but could also be beneficial to snail hosts. Sugar degradation is likely involved in energy generation for the bacteria. Polyamines are linked to bacterial growth and survival by reducing oxidative stress (Murray Stewart *et al.*, 2018) which could be generated by the presence of the hemocytes (Castillo *et al.*, 2020). Production of critical metabolites by the hemolymph microbiome could support sustainable production of these metabolites. While pathway inference using 16S data can give good indication of the pathways present and differentially represented between samples, a more robust approach using metagenomics is required to confirm these inferences.

This microbiome could also be involved in snail defense either directly by controlling pathogens or indirectly by stimulating the snail immune system or competing with pathogens. Snails are susceptible to pathogens ranging from bacteria (Duval *et al.*, 2015) to multicellular parasites (Laidemitt *et al.*, 2017) such as schistosomes causing damages to the snail organs.

### 5. Role of the hemolymph microbiome in the snail-schistosome interactions

We hypothesize that hemolymph microbiome could be a third player in the interaction between the snail and its schistosome parasite. Such tripartite interactions are increasingly recognized to modulate infection outcome and could play a major role in evolution of host-parasite systems (Dheilly, 2014; Ford *et al.*, 2017). The hemolymph is known to be involved in snail defense against schistosomes by the action of hemocytes and humoral factors (Castillo *et al.*, 2020). Snail genetics has also been shown to be a major determinant of snail resistance (Tennessen *et al.*, 2015). However, the microbiome could enhance or reduce snail resistance. Enhancing resistance could occur by direct interaction with the parasite (through competition or parasite infection) or by stimulation of the immune system. The microsporidia *C. owczarzaki* showed direct interaction with S. mansoni sporocysts leading to their death while leaving the snail unharmed (Stibbs *et al.*, 1979; Hertel *et al.*, 2002). These microsporidia were found in higher density in resistant snail populations (Stibbs *et al.*, 1979). This could result from resistant snails actively selecting this particular microorganism. Furthermore, the abundance of bacteria from the *Gemmatimonas* genus in whole snails has been associated with snail genotypes linked to schistosome resistance (Allan *et al.*, 2018), genus which is also present in the hemolymph of our snail populations. While it is unknown if the bacteria associated with the resistant genotype are actually involved in resistance, this showed that genetic control of microbial composition occurs in snails and could influence interactions between snails and schistosomes.

Beyond the snail hemolymph microbiome, we may also have to consider the influence of a potential schistosome microbiome. Role of parasite microbiomes in host-parasite interactions is still poorly understood but could be critical as stated by the Parasite Microbiome Project (Dheilly *et al.*, 2017). Non-human infecting trematodes are known to harbor *Neorickettsia* endosymbionts (Jenkins *et al.*, 2019) and these might also be present in schistosomes (Buddenborg *et al.*, 2017). In addition, schistosome larvae could acquire microorganisms during their free living stages (miracidium and cercariae) or during their development into the snail which could influence the outcome of the infection in snail and mammalian hosts. For instance, the entomopathogenic nematodes *Heterorhabditis* and *Steinernema* rely on their gut bacteria to kill and consume their insect host (Jenkins *et al.*, 2019). The schistosome microbiome, if present, could facilitate the development of developing schistosome larvae inside the snail by disturbing the hemolymph microbiome or by interacting with the snail immune system.

In addition to the fact that *Biomphalaria spp*. snails are intermediate hosts of *S. mansoni*, these snails have several attractive features that make them a promising model system for studying invertebrate-microbiome interactions. Namely (i) they are easy to grow in large numbers in aquaria or plastic trays and experimentally tractable, (ii) they are relatively large and hemolymph can be easily sampled, (iii) as these snails are hermaphrodite, inbred snail lines are easily generated by serial inbreeding to minimize host genetic effects (Bonner *et al.*, 2012), (iv) improving genomic resources and tools for gene silencing are available (Knight *et al.*, 2011; Allan *et al.*, 2017). We anticipate that *Biomphalaria spp*. may join the list of non-traditional model invertebrate species (Douglas, 2019) that are now being exploited for asking both pure and applied questions about microbiome biology.

## EXPERIMENTAL PROCEDURES

### 1) Biomphalaria spp. *hemolymph* collection

*Biomphalaria spp*. freshwater snails are the intermediate host for schistosome trematode parasites. In this study, we sampled only hemolymph (i.e. blood) from schistosome uninfected snails. All our *Biomphalaria spp*. snails were reared in 10-gallon aquaria containing aerated freshwater at 26-28°C on a 12 hour light – 12 hour dark photocycle and fed ad libitum on green leaf lettuce. An advantage of Biomphalaria snails is their large size and their flat shell. This allows easy access to the heart and makes collection of ^~^80-100 μL of hemolymph possible. All snails used in this study had a shell diameter between 10-15 mm.

For each individual snail, we disinfected the shell 3 times with 70% ethanol. We validated the effectiveness of the disinfection method on an independent set of snails. We swabbed shells before and after the 70% ethanol cleaning using sterile polyester tipped swabs (Puritan), extracted DNA and amplified 16S rDNA as described in the next section. This showed no detectable rDNA and confirmed that collected hemolymph was not contaminated with shell microbiome (Supp. Fig. 1). After shell cleaning, we collected the hemolymph by heart puncture using a sterile 1 mL syringe and a sterile 22 gauge needle. The hemolymph collected was immediately placed in a sterile 1.5 mL pre-cooled microtube on ice.

We collected hemolymph from 5 individual snails from each of 7 different populations of *Biomphalaria spp*. (Table 2, Figure 1). We also sampled water from the aquaria where each *Biomphalaria spp*. populations were raised as a control, to characterize the microbiome from the snails’ environment. Immediately after collection, samples were snap-frozen in liquid nitrogen and stored at −80°C until processing. All the samples were collected by a single experimenter, in a 2h window.

### 2) 16S rDNA library preparation and sequencing

We extracted DNA from the snail hemolymph and water samples using the Blood and Tissue kit (Qiagen), following the manufacturer’s protocol, with minor modifications. All the DNA extractions were processed the same day, using a single Blood and Tissue kit (Qiagen) kit, by a single experimenter. We also added blank controls starting at the DNA extraction step to check for potential contaminant coming from the DNA extraction kit and PCR reaction kit. To verify the reproducibility of our process for DNA extraction, library preparation and sequencing, we split each hemolymph or water sample in two aliquots and processed them independently. For each replicate sample, we extracted total DNA using 40 μL of the hemolymph or water samples. We incubated each replicate for 45 minutes at 56°C in a water bath. We recovered final gDNA in 50 μL of elution buffer (Qiagen).

We prepared the 16S rDNA libraries under a biosafety cabinet to avoid contamination, and in triplicate to avoid biases potentially introduced during PCR reaction step. We amplified 250 bp of the 16S rDNA V4 region from bacteria and archaea, using specific primers (515f and 806rB) designed by the Earth Microbiome Project (Apprill *et al.*, 2015). Each of the 515f forward primers were barcoded to allow sample identification. The 16S rDNA V4 reaction mixture consisted of 4 μL of 1X 5 Prime Hot Master Mix (QuantaBio), 2 μL of sterile water, 1 μL of 2 μM 515f barcoded primer, 1 μL of 2 μM 806rB primer (Supp. Table 4), and 2 μL of total gDNA template. Libraries were amplified using the GeneAmp PCR system 9700 (Applied Biosystems) as follows: 95°C for 3 minutes, then 35 cycles of 95°C for 45 seconds, 50°C for 1 minute, 72°C for 90 seconds, then 72°C for 10 minutes. We electrophoresed 2 μL of each PCR product on agarose gel (2%) to check amplicon size, and absence of contamination in the controls (Supp. File 7). We pooled the triplicate PCR products and the 24 μL of final volume were purified by adding 44 μL of AMPure XP beads (Beckman Coulter), mixing with pipette followed by a 5 minute incubation. We separated the beads on a 96 well-plate magnetic device (5 minutes), discarded the cleared solution from each well, and washed beads twice in 200 μL of freshly-prepared 70% ethanol (1 minute), while maintaining the plate on the magnetic device. After drying beads (3 minutes), we eluted each library by adding 20 μL of nuclease-free sterile water, mixing the beads and water by pipetting followed by a 2 minute incubation. Then we separated the beads on the magnetic device (3 minutes) and placed the purified 16S rDNA V4 barcoded libraries in a fresh 96 well-plate. All incubations were done at room temperature. We quantified these libraries using the Picogreen quantification assay (Invitrogen) following the manufacturer’s instructions. Finally, we made an equimass pool of all the libraries using 15 ng of each library. We quantified the final pool fragment size using the 4200 TapeStation (DNA screen tape, Agilent Technology) and molarity using the Kappa library quantification kit for the Illumina sequencing platform (following the manufacturer’s recommendations). We sequenced the pool on an Illumina MiSeq sequencer at Texas Biomedical Research Institute (San Antonio, Texas). Raw sequencing data are accessible from the NCBI Sequence Read Archive under BioProject accession number PRJNA613098.

### 3) Sequence processing

Commands and scripts used for the sequence processing and downstream analysis are available in a Jupyter notebook on Zenodo (https://doi.org/10.5281/zenodo.4020011).

We demultiplexed the raw sequencing data obtained from the MiSeq using bcl2fastq (v2.17.1.14, Illumina) with the default parameters. We used the fastq files generated as input for Qiime2 (v2019.4) (Bolyen *et al.*, 2019). We checked the read quality from the results of the demux module. We then denoised and clustered the sequences into amplicon sequence variants (ASVs) using the denoise-paired command from the dada2 module with a max expected error of 5. We determined the taxonomy of the ASVs using the SILVA database and the classify-consensus-vsearch command of the feature-classifier module using a 97% identity threshold. ASVs with unassigned taxonomy were blasted against the NCBI nt database using megablast from blast+ (v2.7.1) to identify eukaryotic contaminants for removal in downstream analysis. We used a relatively lenient e-value threshold (1e-2) to increase our power of detection. To build a phylogenetic tree, we aligned the ASVs and masked the highly variable positions using the mafft and mask commands from the alignment module. Finally, we built and rooted a tree from the alignment using the fasttree and midpoint-root commands from the phylogeny module.

### 4) Functional inference of the microbiome

We inferred the microbiome’s function using PiCRUST2 software (v2.1.4b). We used the Qiime2 processed data and exported the ASV representative sequences of the taxonomy analysis and the abundance table. We placed the sequences into the reference tree, performed hidden-state prediction of gene families and generated metagenome prediction. We performed pathway-level inference and finally add functional description to the generated files. Finally, we performed a differential pathway abundance using the ALDEx2 R package (v1.16.0) (Fernandes *et al.*, 2014) on the unstratified pathway prediction.

### 5) Estimation of bacterial abundance by quantitative PCR

We performed real time quantitative PCR (qPCR) reactions to estimate the number of bacterial 16S rDNA in each hemolymph and water replicated sample. Reactions were performed in duplicate using the QuantStudio 5 (Applied Biosystems) as follows: 95°C for 10 minutes, then 40 cycles of 95°C for 15 seconds and 60°C for 1 minute. Duplicate reactions showing a difference in C_T_ greater than one were rerun. The reaction mixture consisted of 5 μL of SYBR Green MasterMix (Applied Biosystems), 0.3 μL of 10 μM forward primer (515f without barcode: GTGYCAGCMGCCGCGGTAA) and 10 μM reverse primer (806rB: GGACTACNVGGGTWTCTAAT) amplifying 250 bp of the 16S rDNA V4 region, 3.4 μL of sterile water and 1 μL of total DNA template. We plotted standard curves using seven dilutions of a purified 16S V4 rDNA amplicon (copies.μL^−1^: 1.34×10^1^, 1.34×10^2^, 1.34×10^3^, 1.34×10^4^, 1.34×10^5^, 1.34×10^6^, 1.34×10^7^). We produced the purified 16S V4 rDNA by amplification of the V4 region of *Escherichia coli* XL 10-gold (Agilent) using TaKaRa Taq R001 AM kit (Clonetech) following the manufacturer’s protocol (PCR cycles: 95°C for 10 min, [95°C for 15 s, 50°C for 30 s, 60°C for 60 s] × 35 cycles, 60°C for 10 min). We purified the PCR product using AMPure XP beads (Beckman Coulter) following the manufacturer’s protocol, and quantified it using Qubit dsDNA BR Assay kit (Invitrogen). We determined the number of copies in the PCR product as follows: PCR product length × (average molecular mass of nucleotides (330 g.mol^−1^) × 2 strands) × Avogadro constant. The number of 16S V4 rDNA V4 copies in each sample was estimated according to the standard curve (QuantStudio Design and Analysis Software). The primer pair (515f/806rB) was chosen to allow direct comparison of qPCR values with Illumina MiSeq sequenced amplicons of the same locus. We used the following equation to normalize the qPCR data:

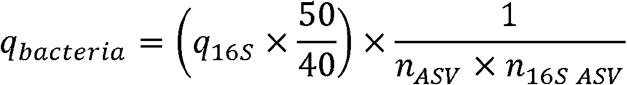

Where the the 16S copy number from the qPCR (q_16S_) was first corrected with the dilution factor between hemolymph and elution volume (40 and 50 μL, respectively) and this value was then normalized using the estimated 16S copy per bacteria from PiCRUST2 (n_16S ASV_) and the abundance of each bacteria from Qiime2 (n_ASV_) (see previous section).

### 6) Statistical analysis

All statistical analyzes and graphs were performed using R software (v3.5.1). We used several packages to compute diversity statistics from the Qiime data: phyloseq (v1.26.1) (McMurdie and Holmes, 2013), microbiome (v1.8.0), and picante (v1.8.1) (Kembel *et al.*, 2010). We compared α-diversity results using a pairwise Wilcoxon test. We used the vegan (v2.5.4) and the pairwiseAdonis (v0.0.1) packages to perform β-dispersion and Permanova tests, respectively. We used the DESeq2 package (v1.22.2) (Love *et al.*, 2014) to identify the differentially represented phyla. We computed centroid distances from the different populations or species using the usedist package (v0.4.0) and tested for significant differences using a t-test or a Wilcoxon test when data did not follow a normal distribution (Shapiro test, p < 0.05).

We compared the microbial density between populations using a Kruskal-Wallis test (data not following normal distribution; Shapiro test, p < 0.05) followed by pairwise comparison post-hoc test. We used Kendall’s tau coefficient to test the correlation between the estimated bacteria density and the observed microbial richness. The confidence interval of significance was set to 95% and *p*-values less than 0.05 were considered as significant.

## ACKNOWLEDGMENTS

We thank Michael S. Blouin from Oregon State University for providing all the inbred *Biomphalaria glabrata* snail lines (Bg26, Bg36 and Bg121), Guillaume Mitta and Benjamin Gourbal (University of Perpignan, France) for providing the BgBRE snails, Philip LoVerde (UT Health, San Antonio) and the Biomedical Research Institute (NIAID Schistosomiasis Resource Center, Rockville, MD) for providing the BgNMRI and BgBS90 snails through NIH-NIAID Contract HHSN272201700014I, and the Theodor Bilharz Research Institute (Giza, Egypt) for providing *B. alexandrina* snails. We thank Shelley Cole and Vanessa Ayala (Texas Biomedical Research Institute) for help with MiSeq sequencing. This research was supported by a Forum grant (14-04491 (FC, WL), a Cowles fellowship (WL) from Texas Biomedical Research Institute and NIH grants (NIH R01AI133749 and R01AI123434 (TJCA)), and was conducted in facilities constructed with support from Research Facilities Improvement Program grant C06 RR013556 from the National Center for Research Resources.

## CONFLICT OF INTEREST

The authors declare no conflict of interest.

## SUPPLEMENTARY MATERIALS

**Supp. Fig. 1 – Shell cleaning effectiveness.** We swabbed snail shells before and after ethanol cleaning and amplified 16S rDNA to test the effectiveness of the cleaning methods. This was done on a snail of each population. L corresponds to the 100 bp ladder; 1 and 2 correspond to swab samples before and after 70% ethanol shell cleaning, respectively; + and − correspond to positive (E. coli DNA) and negative (water only) PCR controls.

**Supp. Fig. 2 – Library rarefaction curve for each snail population.** Curves represent the number of bacterial species identified while subsampling a defined number of 16S V4 sequences. Each curve reached a plateau phase indicating that sampling more sequences did not reveal new bacterial species and therefore that the sequencing was sufficient to capture all the diversity present in the hemolymph sample.

**Supp. Fig. 3 – Correlation of α diversity indices between replicates.** For the six different alpha diversity indices, we performed a correlation analysis between the two replicates and showed that they are highly correlated confirming the reproducibility of our methods. *R* refers to the Pearson’s correlation coefficient (ranging from 0 for no correlation, to 1 for perfect correlation) and *p* refers to the p-value associated.

**Supp. Fig. 4 – Principal coordinates analysis (PCoA) of β diversity indices.** For each index (Jaccard, Unweighted unifrac, Bray, Weighted unifrac), the first three axes of the PCoA are represented in pairwise. The two first indices represent a qualitative measure (absence or presence of bacterial taxa) while the two last are quantitative (adjusted with taxa abundance). A and B refer to replicate 1 and 2, respectively.

**Supp. Fig. 5 – Heatmap of the 94 specific ASVs to snail hemolymph or water microbiomes.** ASVs found in either snails but not water, or in water only, on in specific population of snails.

**Supp. Fig. 6 – Taxonomic diversity in each replicated sample.** This figure is similar to Fig. 4 with the addition of the replicated libraries. Each replicated library clustered together showing the reproducibility of the protocol.

**Supp. Fig. 7 - 16S library preparation gels.** Each library was prepared in triplicate in separated plates. Failed amplifications were run again (plate 4). Gels showed absence of amplification in the negative control (sterile water treated alongside samples).

**Supp. Table 1 – Library statistics.**

**Supp. Table 2 – Differentially represented phyla.**

**Supp. Table 3 – Differentially represented metabolic pathways in snails and water samples.**

**Supp. Table 4 – Earth Microbiome Project primers used for generating each library.**

## Notes

### Competing Interest Statement

The authors have declared no competing interest.

### Summary of Updates

Main updates: * Demonstration of bacterial contamination removal from the snail shell * Unassigned ASV screened for eukaryote contaminants * b-diversity analysis revised

