## Supplementary figures and images for "The hemolymph of *Biomphalaria* snail vectors of schistosomiasis supports a diverse microbiome"

### Supp. Fig. 1

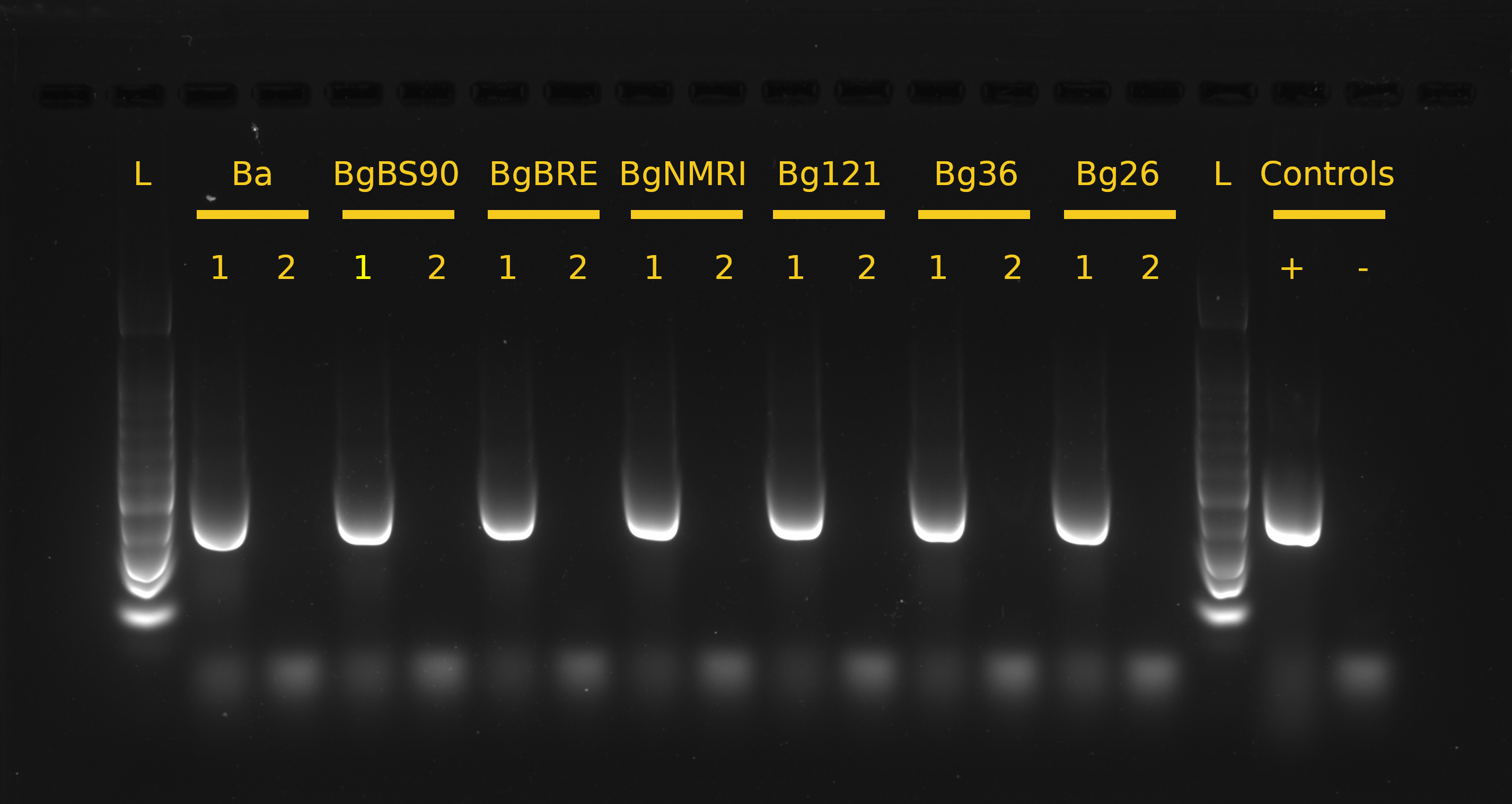

### Supp. Fig. 2

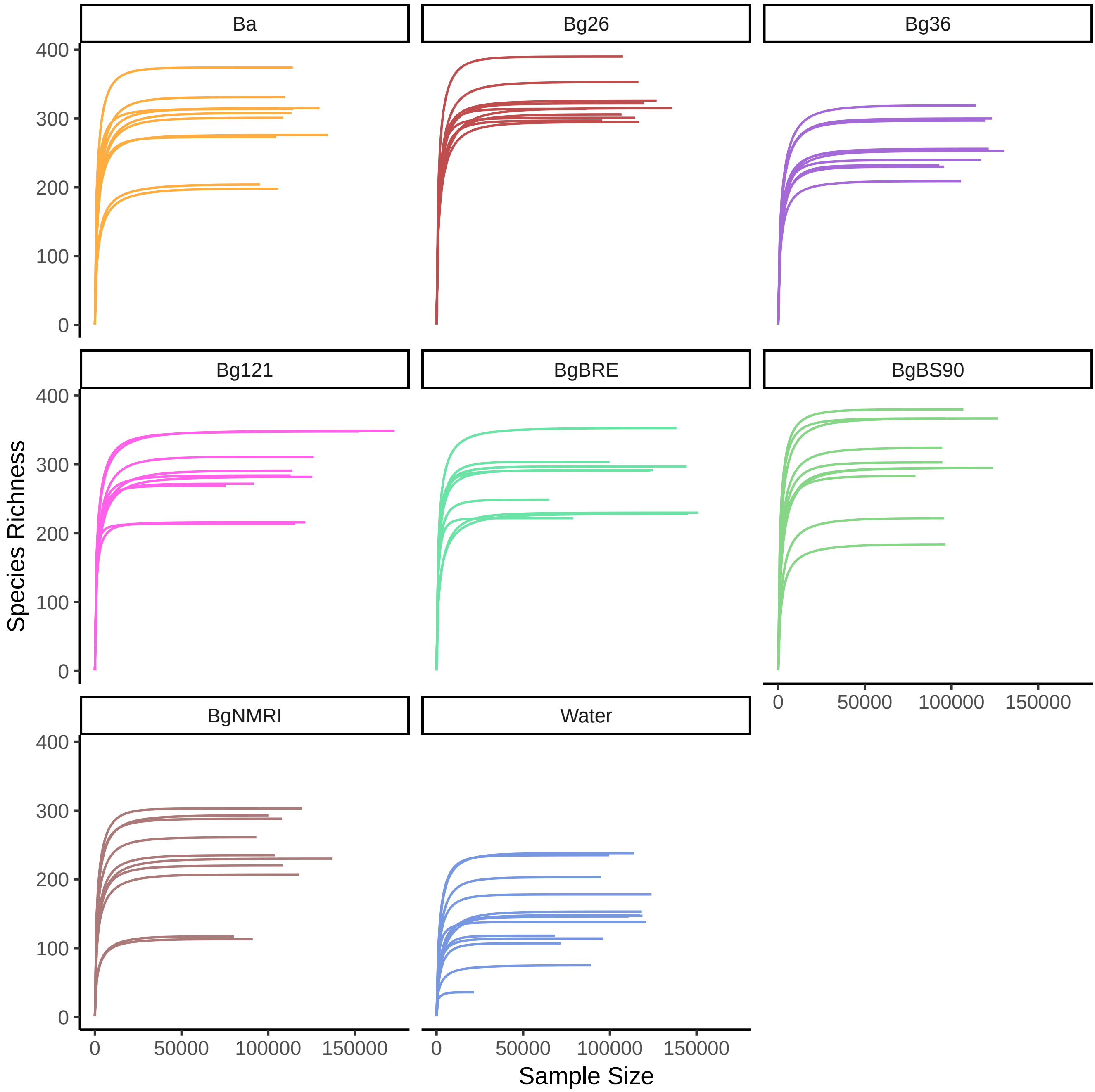

### Supp. Fig. 3

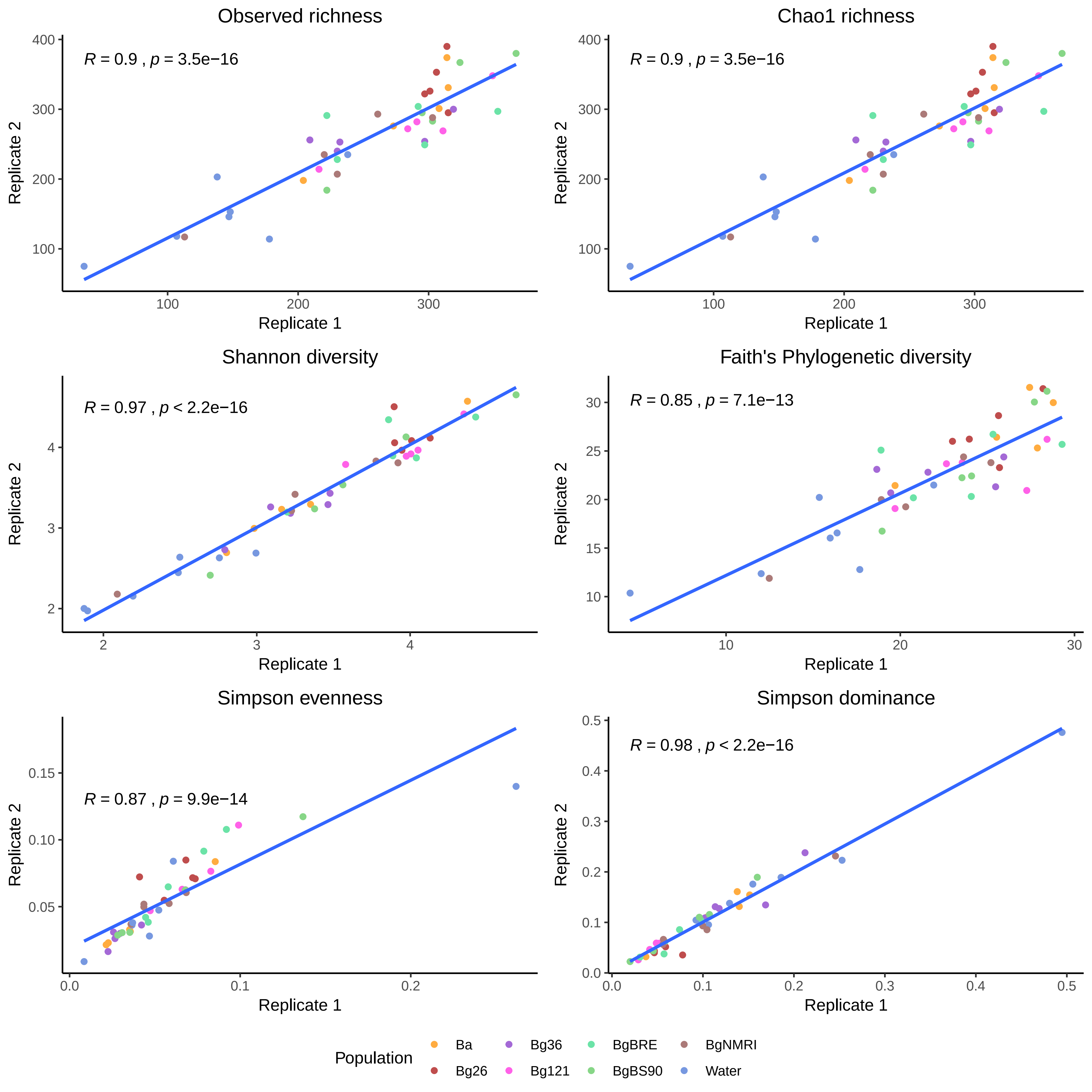

### Supp. Fig. 4

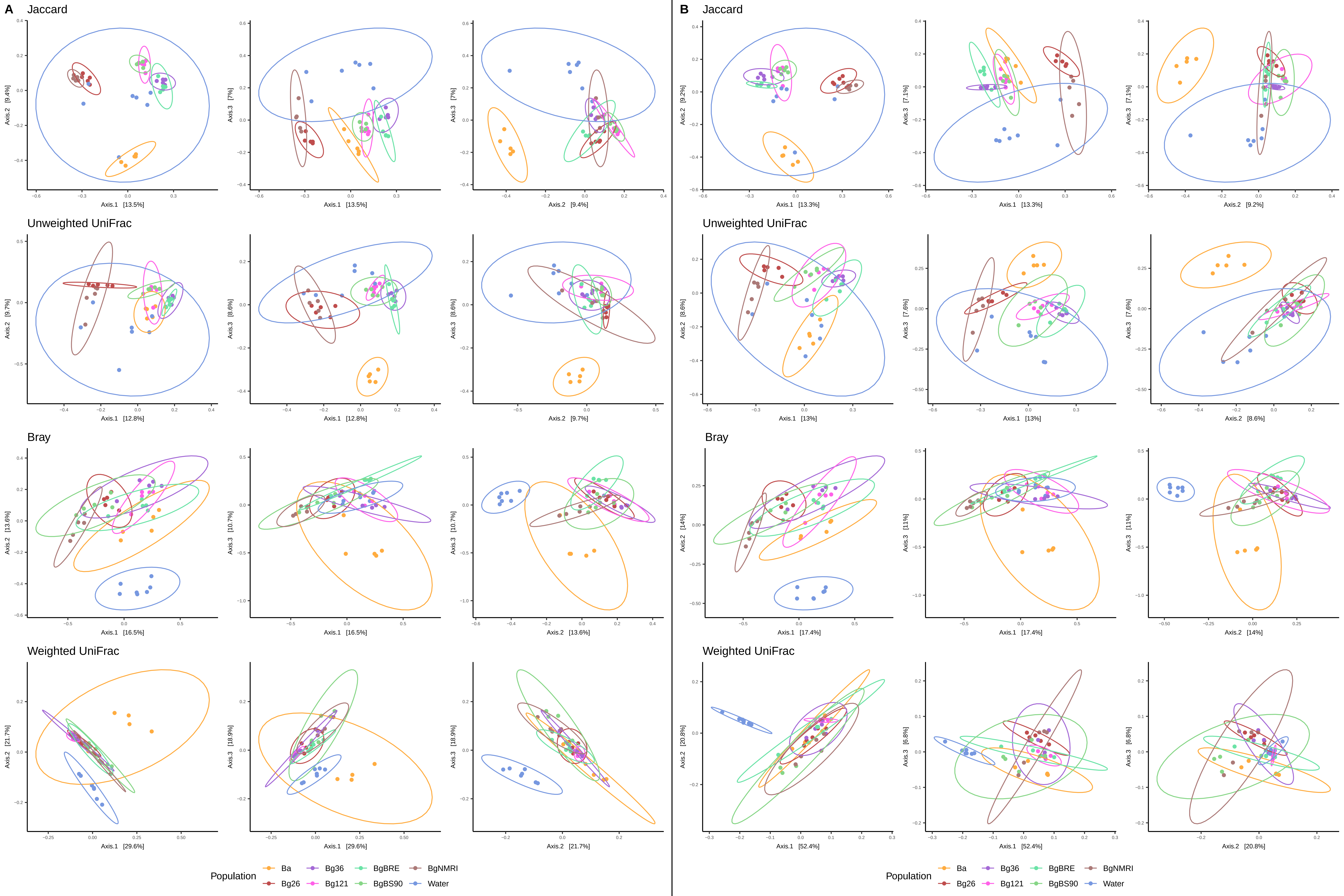

### Supp. Fig. 5

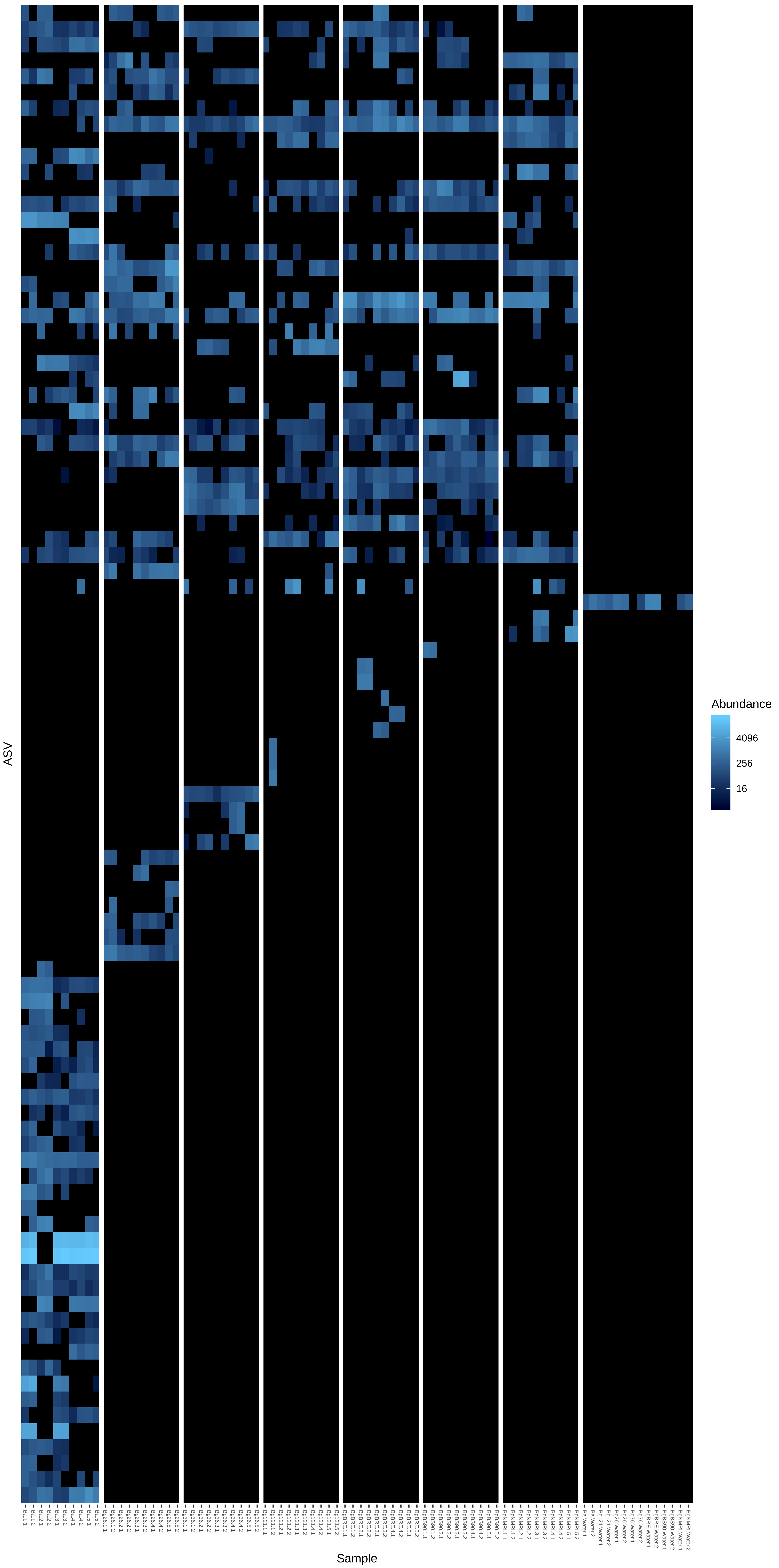

### Supp. Fig. 6

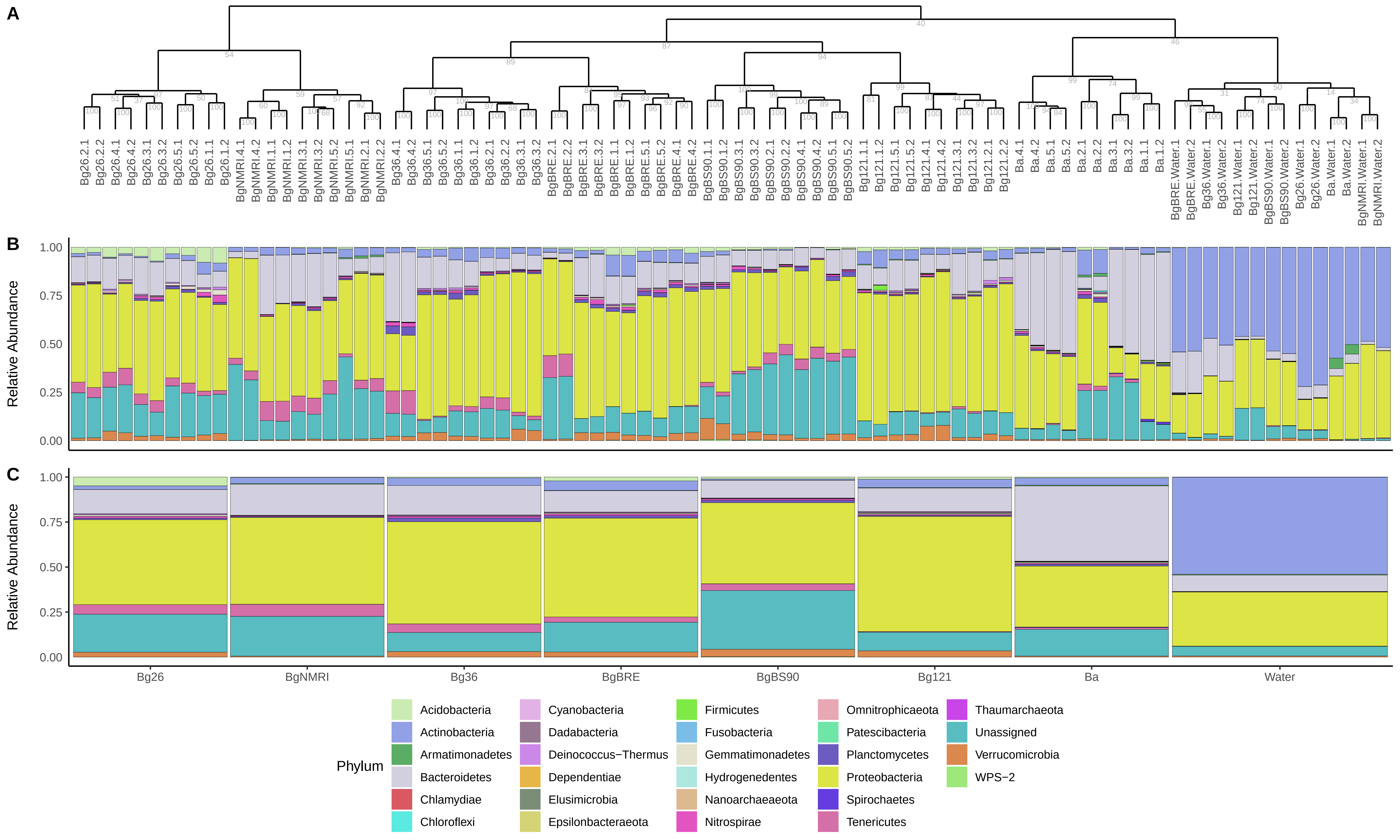
