## Supplementary material for "The hemolymph of *Biomphalaria* snail vectors of schistosomiasis supports a diverse microbiome": Supp. Fig. 7

### 16S library preparation gels

Plate 1
(technical replicate)

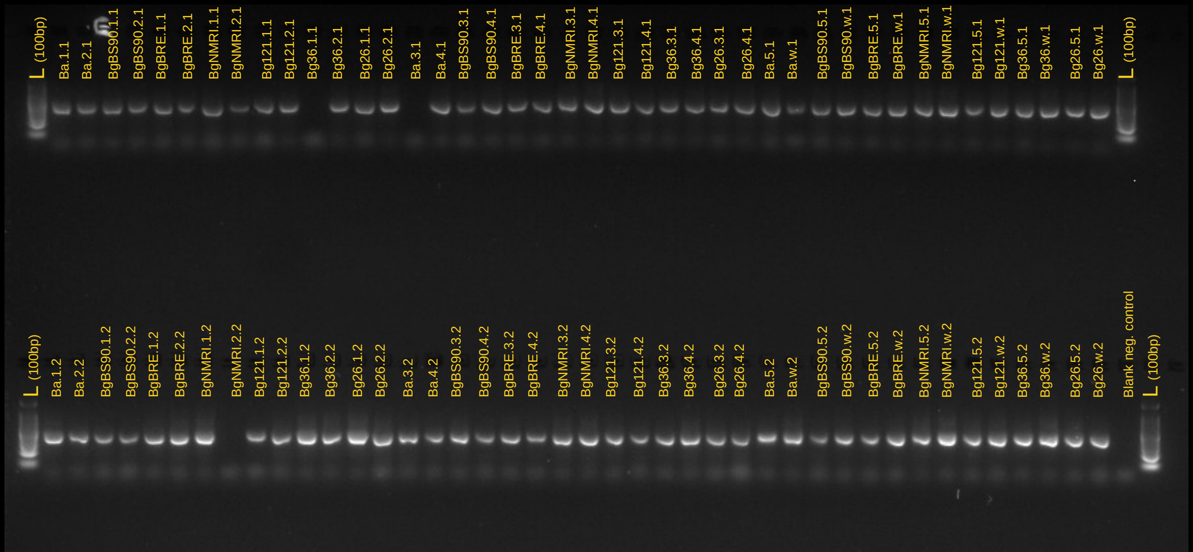


Plate 2
(technical replicate)

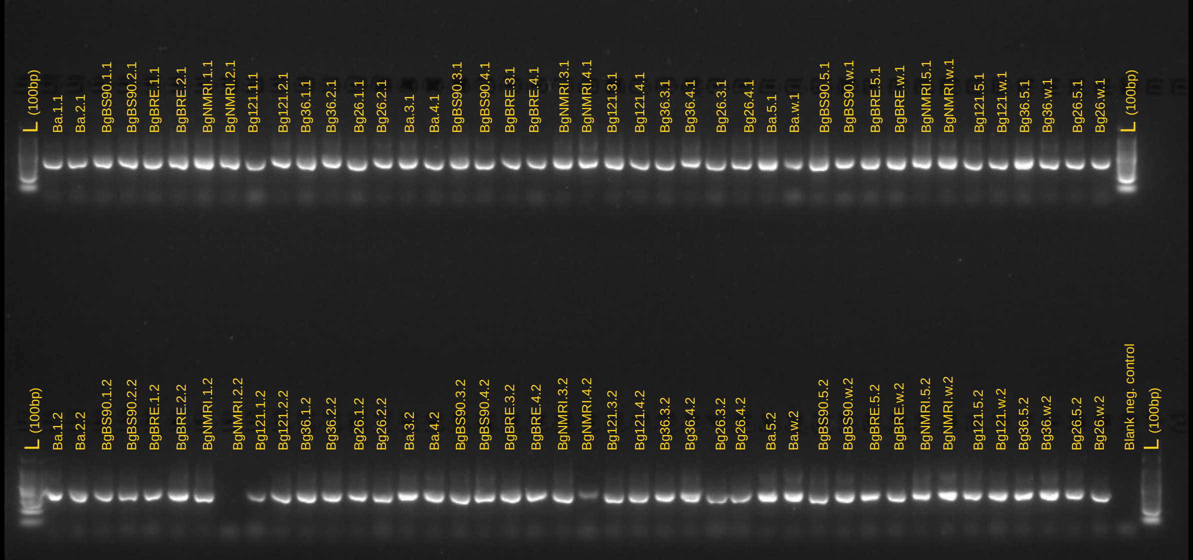


L : Ladder 100bp
w : tank water

Plate 3
(technical replicate)

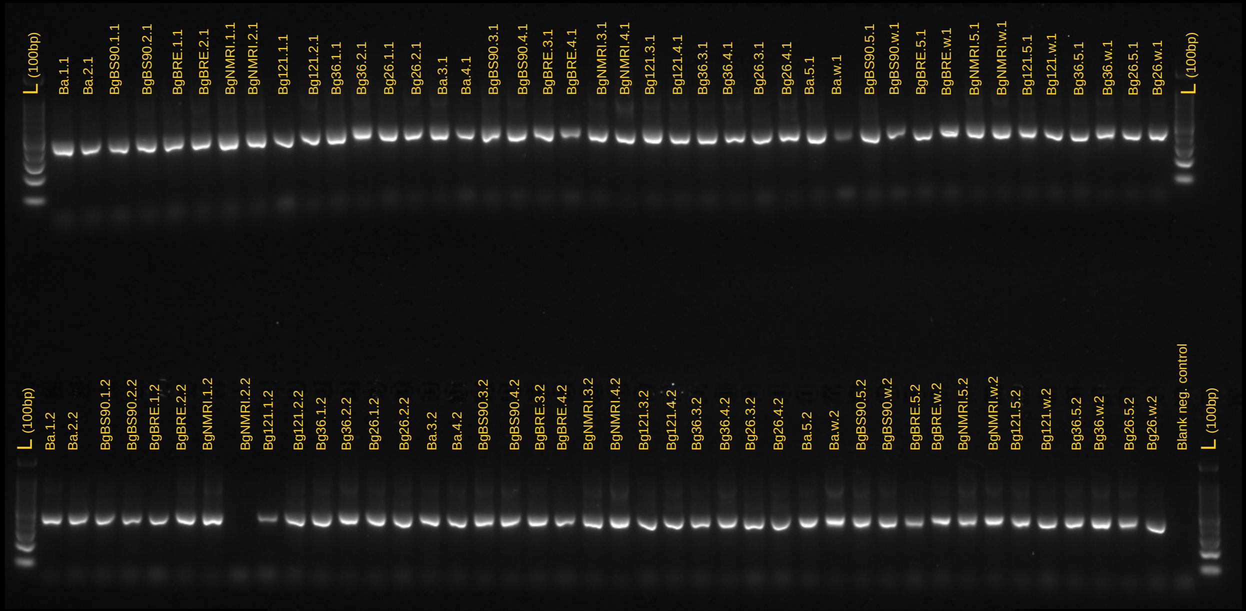


Plate 4
(samples rerun when the first 16S PCR failed)

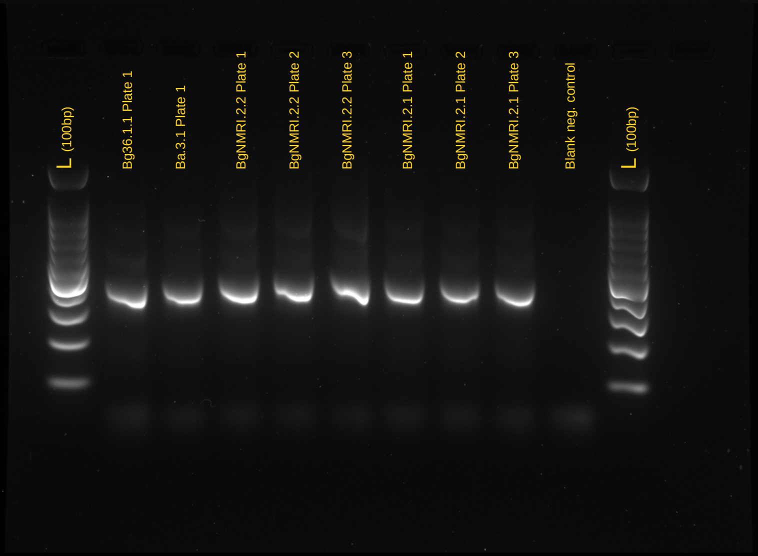


L : Ladder 100bp
w : tank water
